# Draft Genome Sequence of *Modestobacter* sp. strain VKM Ac-2676 from soil in the area of sharply continental climate

**DOI:** 10.1101/179697

**Authors:** Oleg V. Vasilenko, Sergey V. Tarlachkov, Irina P. Starodumova, Elena V. Ariskina, Kseniya I. Ariskina, Alexander N. Avtukh, Lyudmila V. Lysak, Miroslav V. Telkov, Lyudmila I. Evtushenko

## Abstract

A draft genome sequence of a novel actinobacterium, *Modestobacter* sp. strain VKM Ac-2676, was derived using Ion Torrent sequencing technology. The genome size is 3.88 Mb with average G+C content of 73.40%. The genes encoding properties relevant to adaptability of this bacterium to a sharply continental climate zone were revealed.

---

The genus *Modestobacter* (family *Geodermatophilaceae*, phylum “Actinobacteria”) includes six validly described species (LPSN; http://www.bacterio.net) (1). A seventh species has recently been proposed (2). The *Modestobacter* species are aerobic, non-spore-forming, heterotrophic, and able to grow on oligotrophic media. They form orange-red or pink colonies usually turning black with age (2, 3). Members of this genus occur in markedly different ecosystems, including extreme ones (2 - 6).

Strain VKM Ac-2676 was isolated from technogeneous soil collected near Orsk town (Southern Urals, Russia) located in a site characterized by dry and hot summers, with temperatures up to +40°C, and cold winters, with temperatures down to -40°C. Based on the 16S rRNA gene sequence analysis, VKM Ac-2676 is the closest to *M. versicolor* CP153-2^T^ (99.52% sequence identity).

The genome sequencing was performed with a semiconductor genome analyzer Ion Torrent PGM™ (Thermo Fisher Scientific Inc., USA) using 400-bp sequencing kit and 318 v2 chip. A total of 539,682 raw reads were assembled *de novo* into 2,200 contigs (21.0-fold peak coverage) using Newbler 2.9 (454 Life Sciences Corporation, USA). Genomic contigs were annotated using the NCBI Prokaryotic Genomes Automatic Annotation Pipeline (PGAAP, version 3.3) (https://www.ncbi.nlm.nih.gov/genome/annotation_prok). The genome size is 3,884,161 bp with average G+C content of 73.40%. The N50 contig is 2,861 bp, largest contig is 22,534 bp. A total of 2,702 protein-encoding genes (1031 of which are hypothetical proteins), 45 tRNAs, 3 rRNAs and 3 ncRNAs were predicted.

WGS analysis revealed that VKM Ac-2676 harbors genes that encode properties relevant to the adaptability of target bacterium to a sharply continental climate. These include genes involved in the heat shock (*hrc*A, *dna*J and *dna*K) and cold shock (Csp family) responses, and also in the trehalose biosynthesis, which play a crucial role in protecting cells against desiccation. The strain has also genes encoding carbon monoxide dehydrogenase (involved in chemolithotrophic metabolism allowing organisms to use CO as a carbon and energy source) (7) and a putative rhodopsin-like protein genes indicative of its probable photoheterotrophic lifestyle. The genome harbors a *rec*O gene and four copies of *rec*Q DNA helicase that play roles in maintaining genomic stability (8, 9). Strain VKM Ac-2676 also contains genes of multienzyme complex involved in UV resistance, such as *uvr*A and *uvr*B, like that reported for *M. marinus* (NC_017955.1) and *M. caceresii* (NZ_JPMX01000000) (2, 5). The protein sequence of *uvr*C gene of strain VKM Ac-2676 showed high similarity (∼95%) to that of a relevant gene in *M. caceresii* 45-2b^T^, however PGAAP marks this gene as pseudogene.

VKM Ac-2676 formed a sister branch to *M. versicolor* in the *gyr*B gene tree with 92.90% sequence identity, that is in the range of similarities (84.91%–96.31%) found between other species of this genus. The above data suggest that strain VKM Ac-2676 represent a novel species of *Modestobacter*. The study of biochemical and physiological characteristics of this strain is in progress.

**Nucleotide sequence accession number(s)**. This Whole Genome Shotgun project has been deposited at DDBJ/ENA/GenBank under the accession MBFF00000000. The version described in this paper is the first version, MBFF01000000.

## ACKNOWLEDGMENTS

Sequencing was performed at the Postgenomics Research Laboratory, SRI PCM FMBA of Russia. We are thankful to Dr. V.V. Babenko and Dr. E.S. Kostryukova for their kind assistance.

## Funding information.

This work was financially supported by the Russian Federal Agency of Scientific Organizations for fundamental studies at IBPM RAS (topic No. 01201350918).

